# Recombination accelerates adaptation across genetic architectures and demographic histories

**DOI:** 10.64898/2026.09.23.753908

**Authors:** Maria Akopyan, Ellie E. Armstrong, Kieran Samuk

## Abstract

According to classic population genetics theory, recombination shapes the efficacy of natural selection. However, whether variation in recombination rate itself translates into meaningful differences in adaptive capacity remains poorly understood. Here, we used forward-time simulations to examine polygenic adaptation in populations colonizing novel environments. We varied recombination rate eightfold, modeled QTL effect size distributions using three gamma distributions, and compared constant and contracted population sizes. Across all conditions, higher recombination rates modestly accelerated adaptation, reducing the time to reach a new phenotypic optimum by 2–15%. The benefit was substantially greater in constant-size populations than in those that had experienced a contraction, but did not differ across genetic architectures. To understand the mechanism of this acceleration, we tracked linkage disequilibrium among adaptation-related QTL throughout adaptation. We found that across simulations, beneficial alleles were consistently in repulsion phase with one another and in coupling phase with deleterious alleles, both signatures of Hill-Robertson interference. Higher recombination attenuated both forms of interference, indicating that the benefit of recombination operated primarily through relieving interference among QTL. These results implicate interference among linked loci as the primary constraint on polygenic adaptation in this context, and demographic history as a critical modulator of the benefit of recombination.

## Introduction

Recombination is a fundamental evolutionary mechanism that shapes population genetic variation and adaptive trajectories. By shuffling alleles across genetic backgrounds, recombination can promote adaptation by generating novel beneficial haplotypes, breaking down interference among linked loci, and freeing beneficial alleles from associations with deleterious ones (Muller 1964; Hill and Robertson 1966; Felsenstein 1974). Conversely, recombination can impede adaptive evolution by disrupting allelic combinations favored by selection (Altenberg and Feldman 1987; Barton 1995; Feldman et al. 1996). For instance, local adaptation in the face of maladaptive gene flow is often achieved through the clustering of adaptive alleles in genomic regions of low recombination (e.g., chromosomal inversions), which shield locally favored allele combinations from dissociation (Kirkpatrick and Barton 2006; Yeaman and Whitlock 2011; Tigano and Friesen 2016). At the same time, recombination suppression can constrain a population’s capacity to respond to future environmental change, by limiting the formation of new beneficial haplotype combinations and trapping alleles in combinations that may become maladaptive (Roesti et al. 2022; Nickel and Foote 2026). Thus, the effects of recombination on adaptation are complex and context-dependent.

The genetic architecture of traits under selection, including the number of loci and effect sizes of contributing mutations, likely plays a central role in determining how recombination influences adaptive dynamics. Many ecologically important traits are polygenic and can draw on standing genetic variation already present in the population rather than relying solely on *de novo* mutations (Orr and Betancourt 2001; Barrett and Schluter 2008). Because polygenic adaptation requires coordinating subtle frequency shifts across many loci simultaneously, selection acts not on individual alleles but on multilocus haplotypes, which recombination can help assemble (Jain and Stephan 2017; Höllinger et al. 2019). When many loci are under selection at once, however, interference among linked variants can reduce the efficacy of selection on any individual locus (Hill and Robertson 1966; Felsenstein 1974). As such, the rate at which recombination breaks down these associations is a potentially critical determinant of the dynamics of polygenic adaptation. However, it remains unclear the specific conditions under which recombination promotes and interferes with polygenic adaptation, and the relative importance of generating favorable haplotype combinations versus relieving interference among linked loci. Direct tests are needed to disentangle the relative contributions of these phenomena and determine whether the balance between these two mechanisms depends on the degree of polygenicity.

Demographic history is also likely to modulate the effect of recombination on adaptation, in part because the amount of fitness-affecting standing genetic variation available for selection to act on depends directly on demographic history (Charlesworth 2009; Wilson et al. 2014). Populations with large effective sizes maintain more segregating variation, providing recombination with more allelic combinations to work with and making selection more efficient. Population contractions, by contrast, deplete standing variation, and while small populations stand to benefit from higher recombination rates that help generate novel allele combinations (Otto and Barton 2001), amplified genetic drift can overwhelm selection and prevent these benefits from being realized (Wang et al. 2026). Despite the clear potential for both genetic architecture and demographic history to shape how recombination influences adaptation, we lack a systematic understanding of how adaptation proceeds across different genetic architectures and demographic scenarios under varying recombination regimes. While considerable theoretical work has addressed the role of recombination in adaptation, much of it has relied on simplified models involving few loci, and detailed simulation studies exploring these interactions across realistic polygenic architectures and demographic scenarios are lacking (Barton 1995; Otto and Lenormand 2002; Höllinger et al. 2019; Hayward and Sella 2022).

The threespine stickleback provides a powerful model system for exploring these dynamics. Ancestral marine populations have independently given rise to countless freshwater populations across the northern hemisphere, and this repeated adaptive radiation proceeds largely from standing genetic variation (Reid et al. 2021). Strikingly, adaptive alleles are disproportionately clustered in regions of low recombination, implicating recombination rate as an important factor in structuring the genomic basis of adaptation in this system (Roesti et al. 2013; Samuk et al. 2017; Kingman et al. 2021). This makes threespine sticklebacks an ideal foundation for parameterizing forward-time simulations, not to model stickleback evolution *per se*, but to leverage empirically grounded parameters to build intuition about how recombination shapes the rate and outcome of adaptation.

Here, we use forward-time simulations inspired and parameterized by threespine stickleback biology to ask: is adaptation limited by recombination? Specifically, we ask how recombination rate, genetic architecture, and demographic history jointly influence the rate and dynamics of adaptive evolution. We contrast three genetic architectures differing in the distribution of mutational effect sizes across a range of recombination rates, under both constant and contracted population sizes. By tracking how quickly populations adapt to a simulated environmental shift and monitoring population genetic summary statistics over time, we characterize the conditions under which recombination accelerates, constrains, or has little effect on the pace of adaptation.

## Methods

### Simulation design

We used SLiM v5.2 (Haller et al. 2025) to implement forward-time simulations of adaptation to a new environment, modeled after the colonization of freshwater habitats by ancestral marine threespine sticklebacks in post-glacial lakes (Bell and Foster 1994; Thompson 1997; Cresko et al. 2007; Hunt et al. 2008). The general structure of these simulations (Fig. 1) was adaptation to a single phenotypic optimum in a large ancestral population, followed by colonization of a new environment and subsequent adaptation to a new phenotypic optimum. In these simulations, we varied three factors of interest: the distribution of mutational effects (three levels), recombination rate (four levels), and demographic history (two levels), for a total of 24 parameter combinations, each replicated 1,000 times. We simulated populations using the default Wright–Fisher population model in SLiM. A schematic of our model can be found in Figure 1 and specific details of the parameters are described below.

**Figure 1.**
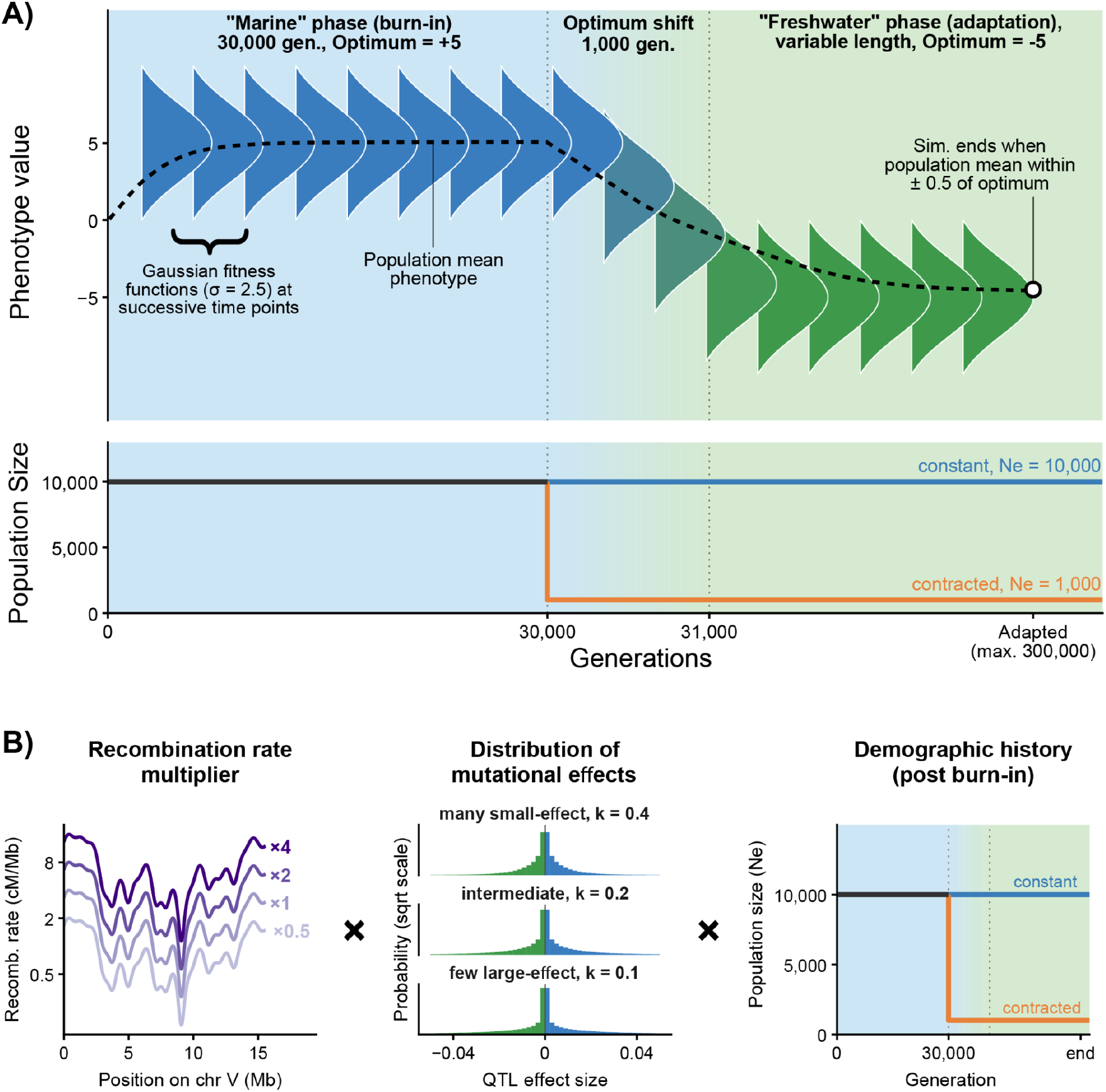
Simulation design overview. **A)** All simulations modeled a two-phase marine-to-freshwater colonization scenario. **B)** Three factors were varied across simulations: recombination rate multiplier, distribution of mutational effects, and post-colonization population size. All 24 parameter combinations were replicated 1,000 times, for a total of 24,000 simulations.

### Genome and recombination

We modeled a single 15.57 Mb chromosome based on stickleback chromosome 5. Recombination rates were drawn from the Puget Sound population map estimated by Shanfelter et al. (2019), averaged into 10 kb windows, and scaled by multipliers of 0.5, 1, 2, and 4. The mutation rate was set to 5 × 10^-9^ per bp per generation (Zhang et al. 2025) across the entire genome. Mutations arising in QTL regions had phenotypic effects drawn from a gamma distribution, while mutations arising elsewhere were neutral. All mutations were modeled as codominant (h=0.5) in their phenotypic effects.

### Genetic architecture

To explore the effect of mutational availability and genetic architecture, we simulated three different distributions of phenotypic (QTL) effect sizes. We modelled these as gamma distributions, parameterized to span a biologically plausible range of architectures, with shape and scale parameters consistent with empirically estimated distributions of fitness effects across animals (Lin et al. 2025). Specifically, all three architectures shared a mean effect size of 0.002 but differed in the shape parameter of the gamma distribution, which controls the degree to which the distribution is heavy-tailed versus evenly spread. Lower shape values produce distributions in which most mutations have very small effects and large-effect mutations are rare. We simulated three distributions with varying degrees of (realized) polygenicity: (1) “many small-effect” (shape=0.4),(2) “intermediate” (shape=0.2); and (3) “few large-effect” (shape=0.1) (Fig. 1). Each mutation arising in the genome was randomly assigned as neutral (probability = 0.98), a positive-effect QTL (probability=0.01), or a negative-effect QTL (probability = 0.01), such that QTL mutations had equal probability of moving the phenotype toward or away from the optimum. Note that by varying the distribution of phenotypic effect of mutations, we allow the specific realized genetic architecture of adaptation to emerge during the simulation rather than specifying it directly, with each mutational effect distribution providing different inputs. This avoids using empirical QTL effect size distributions as the basis of our genetic architectures, as these have well known biases (Rockman 2012). For simplicity, we refer to our different parameterizations of the QTL effect size distributions as “genetic architecture” herein.

### Demographic histories

The two demographic scenarios included either a constant effective population size (*N*_*e*_; hereafter population size) of 10,000 diploid individuals throughout the simulation or a population contraction scenario in which population size was 10,000 during burn-in and reduced to 1,000 following burn-in. While these population sizes are smaller than those observed in many natural stickleback populations, they capture the relevant dynamics of selection and recombination while remaining computationally tractable.

### Selection and adaptation

The phenotype of each individual was calculated as the additive sum of effect sizes across all QTL mutations carried in its genome, and individual fitness was determined by mapping phenotype onto a Gaussian fitness function with a set optimum (see below) and a standard deviation (σ) of 2.5. In our simulated adaptation scenarios, populations evolved first a 30,000-generation burn-in under a phenotypic optimum of 5 (the “marine” phase), after which the optimum shifted linearly over 1,000 generations to a new optimum of −5 (the “freshwater” phase). Simulations ran until the population mean phenotype reached within 0.5 phenotypic units of the new optimum, or for a maximum of 300,000 generations.

### Tracked summary statistics

For each replicate, we recorded the number of generations to adapt, defined as the number of generations at which the population mean phenotype first reached within 0.5 phenotypic units of the new optimum.

Beginning at the end of burn-in and continuing every 2,000 generations through the end of the simulation, we recorded: allele frequencies at all QTL, population mean phenotype, and genome-wide nucleotide diversity (π, calculated using SLiM’s built-in *calcPi* function).

### Distributions of realized QTL effect sizes, selection coefficients, and counts

To characterize the effect-size and selection coefficient distributions of segregating QTL, we subsampled 50 replicates per parameter combination (1,200 total). To assess how selection reshaped the distribution of mutational effects, we compared the realized effect sizes of QTL in these replicates to their input distributions, reconstructing each input distribution from the same gamma mean and shape parameters used in the simulations and matching the realized QTL count per architecture. Because SLiM records phenotypic effect sizes, the effect-size distribution does not directly represent the realized distribution of fitness effects (DFE). We therefore also estimated a selection coefficient (*s*) for each QTL as a proxy for the DFE, taken as the slope of a logistic regression of allele count on generation, where allele count was the frequency multiplied by 2N and rounded to the nearest integer. We estimated *s* only for QTL with at least three timepoints after burn-in that varied in frequency, and truncated each allele-frequency trajectory at the first generation of fixation or loss, since a trajectory that has flatlined no longer informs its rate of change and destabilizes the model fit. We recorded convergence status for each model fit and excluded QTL with non-converging models from downstream analyses. Note that QTL that were lost or fixed rapidly may be underrepresented in these estimates.

Using the same subsampled replicates, we classified the origin of each QTL as standing variation (segregating at the end of burn-in) or *de novo* (arising after burn-in), retaining only QTL with effects directed toward the phenotypic optimum and increasing frequency across the simulation. We tested whether total QTL counts and the relative contribution of standing versus *de novo* variation differed among genetic architectures, demographic histories, and recombination rates using a negative binomial GLM (total counts) and a binomial GLM (proportion standing), with all predictors and their interactions included. Significance of model terms was assessed using Type II likelihood ratio tests, and pairwise contrasts were estimated to characterize the direction and magnitude of significant effects. Additionally, we tested whether recombination rate multipliers affected the effect-size and selection-coefficient distributions using

Kruskal-Wallis tests. We then used two- and k-sample Anderson-Darling tests to compare realized effect-size distributions to their corresponding input distributions, and to test for differences in effect-size and selection-coefficient distributions among genetic architectures and demographic histories.

### Adaptation rate across recombination rate multipliers, demographic histories, and genetic architectures

Adaptation rate was defined as the number of generations after burn-in required for the population’s mean phenotype to reach the new optimum (–5 ± 0.5). We tested whether adaptation rate varied with respect to recombination rate multiplier, demographic history, and genetic architecture using a three-way ANOVA on log-transformed generations to adapt, with all factors and their interactions as fixed effects. We obtained estimated marginal means for each parameter combination, back-transformed to the response scale.

Because recombination rate multipliers were equally spaced on a log2 scale (0.5, 1, 2, and 4), we used orthogonal polynomial contrasts to test for linear and quadratic trends in adaptation rate across recombination rate multipliers within each demographic history and genetic architecture combination, with p-values adjusted using the false discovery rate (FDR) method.

### Population phenotype and nucleotide diversity trajectories

We summarized each replicate’s initial phenotypic and diversity dynamics as the slope of a linear regression against generation over the first 4,000 generations following burn-in. We additionally quantified diversity recovery as (π_final_ − π_min_)/(π_initial_ − π_min_), where π_initial_ is diversity at generation 30,000, π_min_ is the minimum diversity observed across the full trajectory, and π_final_ is diversity at the last tracked generation, a measure that does not depend on trajectory length. We compared each of these three metrics, initial phenotypic rate of change, initial diversity decline rate, and diversity recovery, across population size history, genetic architecture, and recombination rate multiplier using Kruskal-Wallis tests, with epsilon-squared (ε^2^) as a measure of effect size.

### Linkage disequilibrium dynamics

To examine patterns of linkage disequilibrium among QTL, we calculated mean linkage disequilibrium (D, using SLiM’s built-in calcLD_D function) among QTL pairs segregating above 5% frequency in a subset of simulations: constant-size populations under the small-effect DFE, with 100 replicates per recombination rate multiplier. LD was calculated separately for beneficial-beneficial, deleterious-deleterious, and beneficial-deleterious QTL pairs, beginning at the end of the burn-in period and recorded every 2,000 generations thereafter. Because simulation length varied across replicates, analyses were restricted to the first 14,000 generations following burn-in, ensuring all replicates were represented at each timepoint. To test the effects of recombination rate on LD, we fit separate linear mixed models for each QTL pair category with recombination rate multiplier (as a factor) and scaled generation as fixed effects and replicate as a random effect. Linear mixed models were fit twice for each QTL pair category: without an intercept to test whether mean D differed from zero within each recombination treatment, and with the 0.5× treatment as the reference level to test whether higher recombination rates significantly altered the magnitude of LD.

Data were formatted, visualized, and fit to statistical models in R v4.4.2 (Team 2024) using the tidyverse (Wickham et al. 2019), MASS (Ripley and Venables 2009), car (Fox et al. 2001), emmeans (Lenth and Piaskowski 2017), rstatix (Kassambara 2019), lme4 (Bates et al. 2015), lmerTest (Kuznetsova et al. 2017), and kSamples (Scholz and Zhu 2025) packages.

## Results

### Population contractions reduce the contribution of standing genetic variation

Genetic architecture strongly influenced the total number of QTL (likelihood ratio test: χ^2^=52647, df=2, p<0.001), with the small-effect architecture yielding more QTL, as expected (Fig. 2A). Demographic history also significantly affected total QTL counts (χ^2^=474, df=1, p<0.001), with constant-size populations yielding more total QTL than contracted populations, a difference that became more pronounced at higher recombination rate multipliers (χ^2^=195, df=3, p<0.001). The relative contribution of standing variation versus *de novo* mutations differed among both genetic architecture (χ^2^=14461, df=2, p<0.001) and demographic history (χ^2^=3628, df=1, p<0.001). The small-effect architecture relied more heavily on *de novo* mutations, though this is, at least in part, because adaptation under the small-effect architecture took longer (described below), providing more time for *de novo* mutations to accumulate. Additionally, constant-size populations drew more heavily on standing variation than contracted populations (χ^2^=3628, df=1, p<0.001) regardless of recombination rate (χ^2^=5.5, df=3, p=0.14), with the effect being more pronounced in the small-effect architecture (Fig. 2A, χ^2^=256, df=2, p<0.001).

**Figure 2.**
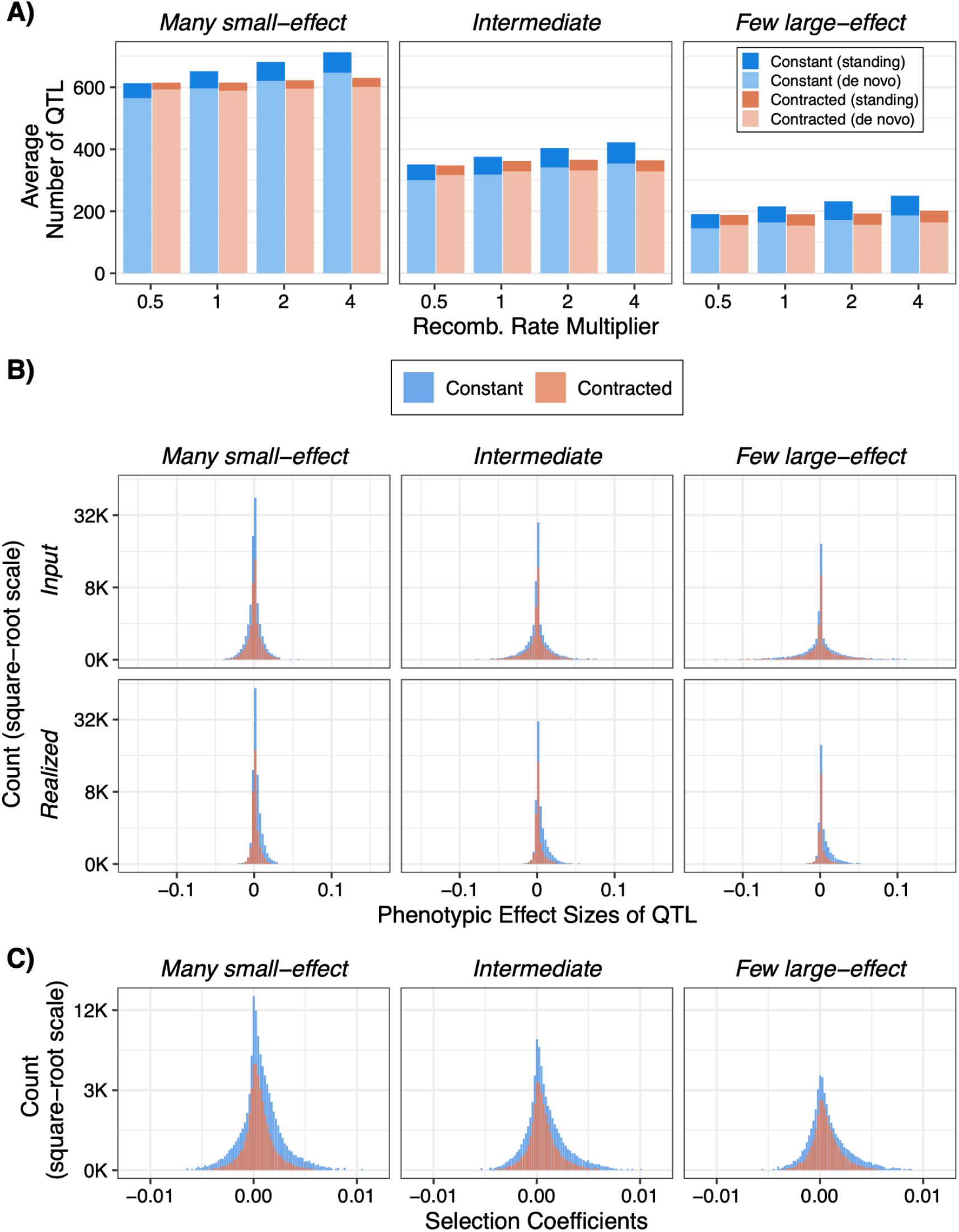
QTL counts, effect sizes, and selection coefficients under three genetic architectures. **A)** Average number of QTL across replicates that were present in the population at the end of burn-in (i.e., standing variation) or arose as *de novo* mutations after the burn-in period. **B)** QTL phenotypic effect sizes representing the input distribution (top) and the realized distribution after adaptation (bottom). Note that the x-axis is reversed so that beneficial effects (negative, toward the phenotypic optimum) appear on the right. **C)** QTL selection coefficients estimated by fitting a logistic regression to each QTL’s allele frequency trajectory. Data shown for 50 randomly sampled replicates per parameter combination (1,200 replicates total).

### Population contractions reduce the efficacy of selection

We compared distributions of phenotypic effect sizes and selection coefficients across parameter combinations. Recombination rate multipliers had a statistically significant but biologically negligible effect on the realized effect-size and selection-coefficient distributions (Kruskal-Wallis ε^2^≤0.0014 and ε^2^≤0.0038 [effect size], respectively). We therefore pooled recombination rate multipliers across levels in subsequent comparisons. Selection significantly reshaped the realized phenotypic effect-size distribution relative to the input distribution in every combination of genetic architecture and population size, such that QTL with effects in the direction of the new optimum were more common than those with opposing effects. This effect was stronger in constant-size populations, consistent with reduced selection efficiency under drift in contracted populations (Fig. 2B). Realized selection coefficients likewise differed significantly by QTL mutation effect size distribution and population size (Anderson-Darling tests, all p<1×10^-16^), with QTL effect size differences again more pronounced in constant-size than contracted populations (Fig. 2C). In particular, in constant-size populations, the median selection coefficient was highest in the small-effect QTL distribution and declined toward the large-effect distribution, while upper-tail coefficients remained similarly elevated.

### Increasing recombination rate accelerates adaptation

Across all six demography and architecture combinations examined, increasing recombination rate significantly reduced the number of generations required for populations to reach the new phenotypic optimum (F_3,23976_=1.00×10^3^, p<0.0001; polynomial contrasts, linear term: all p<0.0001 after FDR correction in every combination), with no reversal of this trend observed under any condition (Fig. 3). Higher recombination therefore consistently facilitated adaptation across the full range of scenarios explored here, though this effect was modest relative to demographic history and QTL effect size distribution.

**Figure 3.**
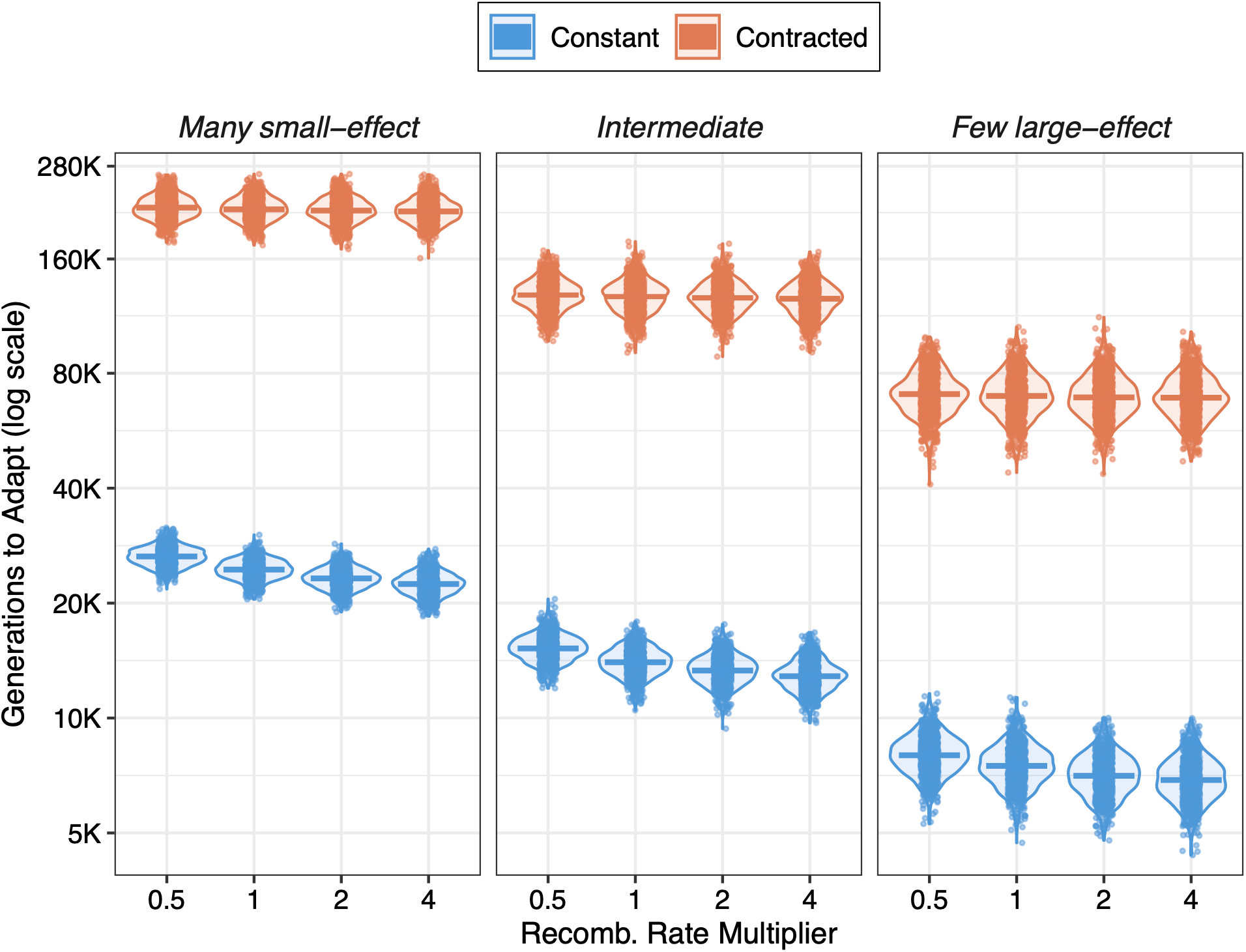
The effect of recombination rate on adaptation speed. Generations (log scale) to reach the phenotypic optimum across recombination rate multipliers under constant (blue) and contracted (orange) population sizes, faceted by genetic architecture. Violins show the distribution across replicates; points show individual replicates; horizontal bars show the mean.

### Demographic history and genetic architecture shape adaptation rate

Demographic history had a much larger effect than recombination rate on adaptation rate (F_1,23976_=3.04×106, p<0.0001), accounting for the large majority of variance explained by the model (83.9% of total sum of squares). Populations that experienced a post-burn-in contraction in population size took roughly 8-to 9-fold longer to adapt than populations that maintained a constant size throughout (e.g., under the small-effect architecture at the lowest recombination rate: 218,057 vs. 26,495 generations), reflecting the reduced efficiency of selection relative to drift, and the smaller effective mutational input, in contracted populations.

Genetic architecture also strongly shaped adaptation rate (F_2,23976_=2.77×10^5^, p<0.0001; 15.3% of total sum of squares): populations adapted faster under architectures with few large-effect loci than under architectures relying on many small-effect loci (e.g., under constant population size at the lowest recombination rate: 7,988 vs. 26,495 generations). This is consistent with expectations that a small number of large phenotypic steps can close a fitness gap more quickly than the sequential substitution of many minor-effect alleles, even when the two architectures share the same mean mutational effect size.

Demographic history’s effect on adaptation speed also depended on genetic architecture (size × dfe: F_2,23976_=184.5, p<0.0001): the slowdown associated with a population contraction increased from 8.2-fold under the small-effect architecture to 8.8-fold under the large-effect architecture.

### Recombination’s benefit depends on demographic history, not genetic architecture

Although recombination consistently accelerated adaptation, the magnitude of this benefit depended on demographic history (recombination rate × demographic history: F_3,23976_=564.6, p<0.0001). Under constant population size, increasing recombination rate from 0.5–4× the empirical map reduced the number of generations to adapt by ∼14–15% within each genetic architecture. In terms of numbers of generations required to reach the new optimum, populations with a 0.5× versus 4× recombination rate adapted in 26,495 versus 22,441 generations under a small-effect architecture (4,054 generation difference, 15.3% reduction), 15,213 versus 12,868 generations under an intermediate-effect architecture (2,345 generation difference, 15.4% reduction), and 7,988 versus 6,882 generations under a large-effect architecture (1,106 generation difference, 13.8% reduction). In contracted populations, increasing the recombination rate multiplier from 0.5× to 4× reduced mean adaptation time by between 1,517 and 4,898 generations, depending on genetic architecture. While these reductions were similar in magnitude to those observed in constant-size populations, because adaptation in contracted populations took substantially longer overall (mean adaptation time between 69,033 and 218,057 generations), the same number of generations amounted to a proportional reduction of only 2.2–2.4%. Polynomial contrasts further showed that the shape of this relationship differed between demographic scenarios: under constant population size, the decline in adaptation time with recombination rate exhibited significant negative curvature (quadratic contrasts: all p<0.0001), indicating a nonlinear relationship with diminishing returns at higher recombination rates, whereas in contracted populations, the relationship did not depart significantly from linear (quadratic contrasts, p≥0.20). This suggests that recombination’s contribution to adaptation is most consequential when population size remains large.

In contrast, recombination’s benefit did not depend on genetic architecture: neither the recombination rate × genetic architecture interaction (F_6,23976_=1.40, p=0.212) nor the three-way interaction among recombination rate, genetic architecture, and demographic history (F_6,23976_=1.23, p=0.289) was significant. The proportional benefit of increased recombination was similar across all three architectures within a given demographic scenario (e.g., under constant population size, a 0.5× to 4× increase in recombination reduced adaptation time by 15.3%, 15.4%, and 13.8% under the small, moderate, and large effect architectures, respectively). Recombination’s effect on adaptation speed was therefore robust to the genetic architecture underlying the trait, at least across the range of architectures examined here.

### Population size shapes phenotype and diversity dynamics

Phenotype trajectories (Fig.4A) recapitulated the pattern observed for adaptation time (Fig. 3), with the initial rate of phenotypic change toward the optimum shaped most strongly by population size history (Kruskal-Wallis ε^2^=0.75), followed by genetic architecture (ε^2^=0.21), and only a small effect of recombination rate multiplier (ε^2^=0.005).

Nucleotide diversity trajectories (Fig. 4B) showed a decline following the initial optimum shift in every replicate, consistent with selection at QTL reducing diversity at linked sites genome-wide. Population size history had the largest effect on the severity of this decline (initial decline rate: ε^2^=0.63; diversity recovery: ε^2^=0.71), with contracted populations showing steeper drops reflecting both the genome-wide effects of linked selection and the direct reduction in diversity associated with the population contraction itself.

**Figure 4.**
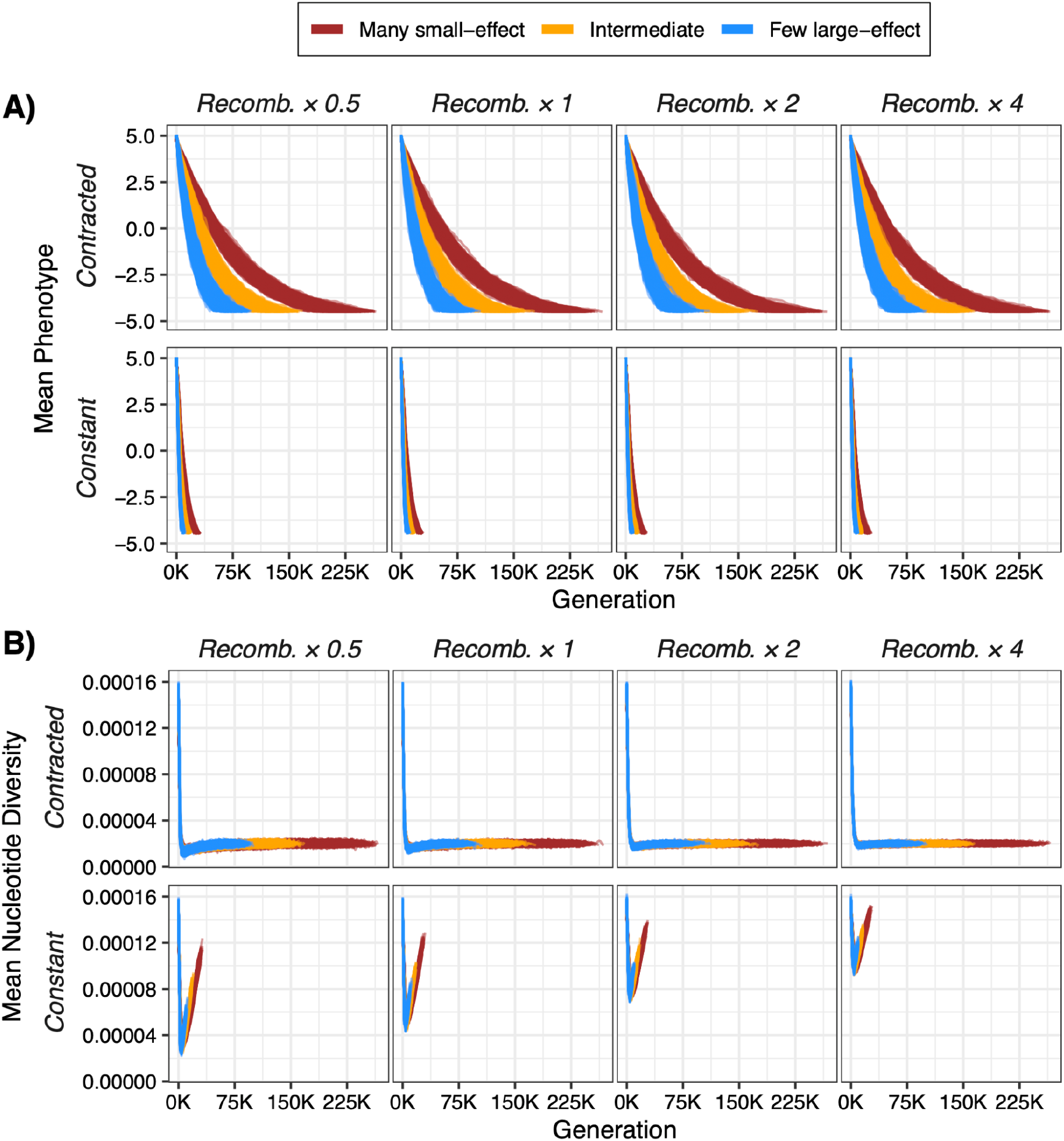
Temporal dynamics of mean phenotype and nucleotide diversity during adaptation. **A)** Mean population phenotype and **B)** mean nucleotide diversity (π) tracked every 2,000 generations across recombination multipliers (columns), demographic histories (rows), and genetic architectures (colors).

Recombination rate multiplier also had a significant effect on the steepness of the initial decline, as did genetic architecture (ε^2^=0.12 for each), consistent with recombination limiting the reach of hitchhiking around selected QTL, though its influence on diversity recovery was comparatively small (ε^2^=0.04).

### Increasing recombination rate accelerates adaptation by reducing Hill-Robertson interference

Mean LD (D) differed in sign between beneficial-beneficial and beneficial-deleterious QTL pairs and was modulated by recombination rate (Fig. 5). Beneficial-beneficial QTL pairs exhibited significant negative LD across all recombination treatments (all t1027≤−5.30, p<0.001), indicating that beneficial alleles consistently resided on different haplotypes (repulsion phase). This pattern is consistent with Hill-Robertson interference, where competing beneficial alleles on separate haplotypes reduce the efficiency of selection and slow the rate of adaptation. Higher recombination rates significantly reduced the magnitude of repulsion relative to the 0.5× treatment (all t3093≥9.26, p<0.001).

**Figure 5.**
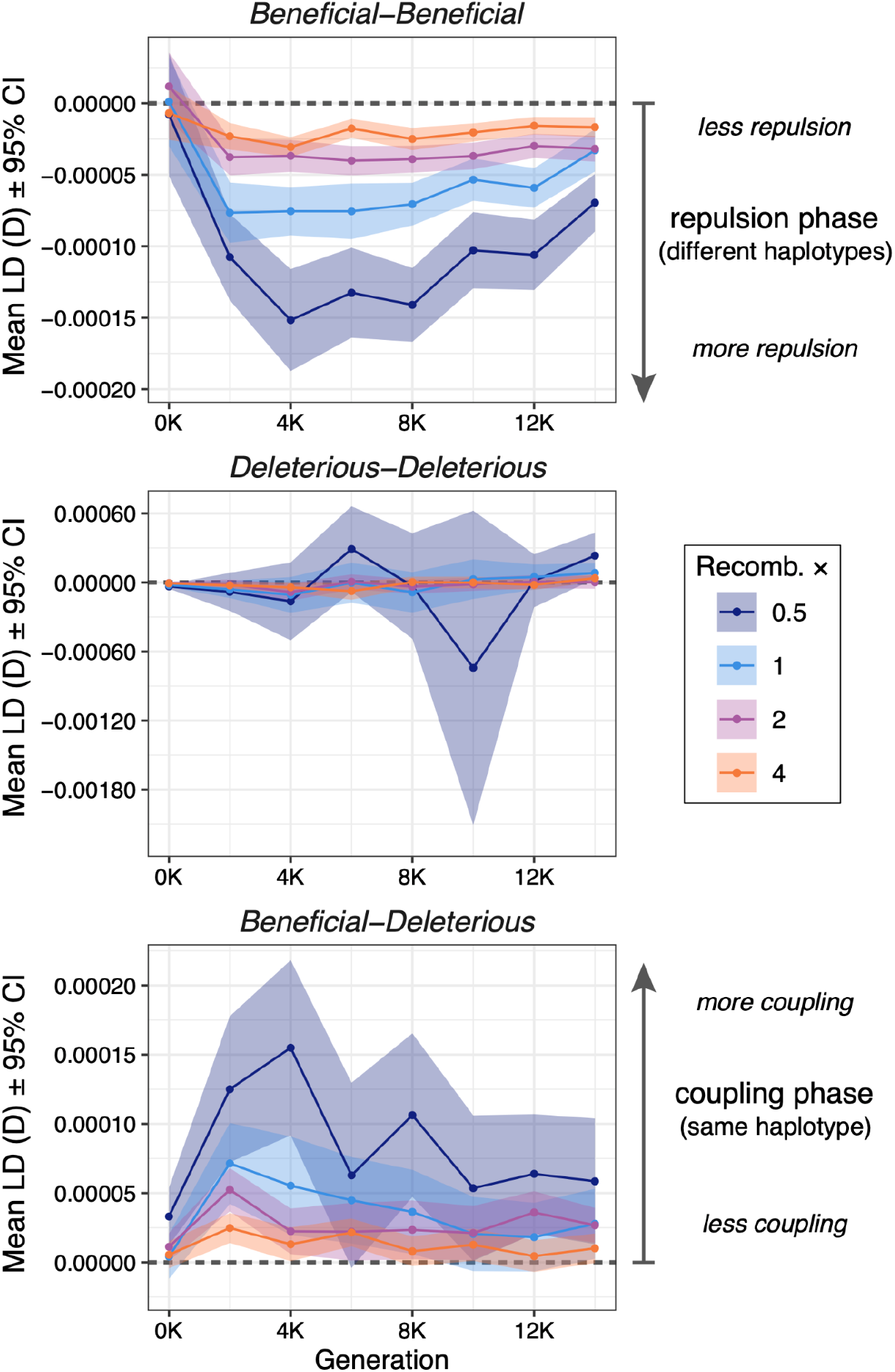
Temporal dynamics of linkage disequilibrium during adaptation. Mean LD (coefficient D) ± 95% confidence intervals across the first 14,000 generations following burn-in for 100 replicates per recombination rate multiplier (colored lines). Panels show QTL pairs grouped by the sign of each allele’s effect on the trait. The dashed line indicates linkage equilibrium (D = 0); negative values indicate repulsion and positive values indicate coupling. Results shown for constant population size and the many small-effect architecture only. Note that the y-axis values vary across panels.

Beneficial-deleterious QTL pairs showed the opposite pattern, with significant positive LD across all recombination treatments (all t940≥2.17, p<0.05), indicating that beneficial and deleterious alleles consistently co-occurred on the same haplotype (coupling phase). This pattern also reflects Hill-Robertson interference, with deleterious alleles hitchhiking with linked beneficial alleles and similarly impeding adaptation. Higher recombination rates significantly reduced the magnitude of coupling relative to the 0.5× treatment (all t3093≤−6.00, p<0.001). Deleterious-deleterious QTL pairs maintained mean D near zero (i.e., linkage equilibrium) across all recombination treatments (all t630≤1.16, p>0.2), and variance was substantially elevated under the 0.5× treatment (Fig. 5). Together, these results suggest that increasing recombination rate accelerates adaptation by reducing Hill-Robertson interference, both between beneficial QTL pairs (reduced repulsion) and between beneficial and deleterious QTL pairs (reduced coupling).

## Discussion

Using forward-time evolutionary simulations parameterized by threespine stickleback biology, we asked whether adaptation is limited by recombination. Across all parameter combinations we examined, higher recombination rates accelerated adaptation. Genetic architecture and demographic history also strongly shaped the rate of adaptation, with less polygenic architectures enabling faster adaptation and population contractions dramatically slowing adaptation. However, these factors did not influence the effect of recombination rate in the same way: higher recombination rates had a greater effect on the rate of adaptation in populations maintained at constant size than in populations that experienced a contraction, but their effect was similar across genetic architectures. Below we discuss the mechanisms underlying these patterns and their implications for understanding when and how recombination facilitates adaptive evolution.

Theory predicts that recombination can either facilitate or impede adaptation depending on the circumstances (Barton and Charlesworth 1998; Otto and Lenormand 2002; Ritz et al. 2017). Across the scenarios we modeled, we found no conditions under which higher recombination slowed adaptation. Increasing the recombination rate multiplier from 0.5× to 4× reduced the number of generations required to reach the new phenotypic optimum under every genetic architecture and both demographic scenarios, by between 2% and 15%. Recombination rates vary tremendously across genomes, and among individuals, populations, and species (Stapley et al. 2017), yet whether this variation translates into meaningful differences in adaptive capacity, that is, whether adaptation is recombination-limited, remains an open question. The eightfold range of recombination rates that we modeled falls well within this variation, indicating that differences in recombination rate of a magnitude commonly observed in nature can affect the rate of adaptation. However, the benefit of increased recombination did not accrue steadily in all cases. In constant-size populations, the decline in adaptation time with increasing recombination rate showed significant negative curvature, indicating diminishing returns at higher rates, whereas in contracted populations, the relationship did not depart from linearity, though recombination’s relative contribution to faster adaptation was considerably smaller. The capacity for recombination to accelerate adaptation is therefore greatest when recombination rates change from low to high in populations that retain substantial genetic variation.

Our LD analyses offer a mechanistic account of how recombination accelerated adaptation in our simulations. Beneficial QTL alleles were consistently found in repulsion phase, indicating that competing beneficial alleles occupied different haplotypes, while beneficial and deleterious alleles were in coupling phase, indicating that deleterious variants hitchhiked with linked beneficial ones on the same haplotype. Both patterns are signatures of Hill-Robertson interference, in which linkage among selected sites reduces the efficacy of selection at each (Hill and Robertson 1966; Felsenstein 1974). Higher recombination significantly attenuated both forms of interference, consistent with the interpretation that increased recombination accelerated adaptation primarily by reducing interference rather than by generating novel favorable haplotype combinations. This result aligns with theoretical predictions that recombination should be most beneficial when multiple selected sites compete for fixation (Hill and Robertson 1966; Otto and Barton 1997). This also explains why recombination’s benefit diminished following a population contraction: a contraction removes segregating variation directly, reduces the input of new mutations, and lowers the efficacy with which selection acts on the variation that remains (Barrett and Schluter 2008; Charlesworth 2009). Recombination can rearrange the variation present in a population, but it cannot create it. Along with matching theory, a key take-away from our simulations is that the general predictions of analytical and more genetically simple models are upheld under complex demography, genetic architectures, and genome-scale variation in recombination rate. This need not have been the case, and demonstrates that classic theoretical expectations are resilient to biological complexity, and can likely be safely extended to real biological populations.

Our finding that recombination consistently accelerated the rate of adaptation raises a question: if higher recombination is advantageous, why do natural populations exhibit substantial variation in recombination rate rather than trending toward uniformly elevated rates? The answer lies in the interplay between direct and indirect selection on recombination, both of which can act in opposing directions (Otto and Lenormand 2002; Drury et al. 2023). Direct selection on recombination stems from its role in chromosome segregation: too few crossovers can produce aneuploid gametes, placing a lower bound on the rate (Hassold and Hunt 2001; Jones and Franklin 2006), while too many increase the risk of ectopic recombination, generating deleterious chromosomal rearrangements (Montgomery et al. 1991). Indirect selection, which operates through recombination’s effects on genotypic variability among offspring (Barton 1995; Otto and Lenormand 2002), is similarly context-dependent. When populations are adapting to novel environments, elevated recombination may be favored because it generates novel allele combinations among offspring. This benefit can arise by bringing together beneficial alleles from different haplotypes and by relieving Hill-Robertson interference among competing selected sites (Muller 1964; Hill and Robertson 1966; Otto and Barton 1997), the latter being the mechanism we confirm here.

Under more stable conditions, however, recombination can disrupt allele combinations assembled by prior selection, imposing a fitness cost that exceeds its benefits, a dynamic known as the Reduction Principle (Otto and Lenormand 2002). Local suppression of recombination can also be favored, playing a central role in preserving locally-adapted haplotypes against maladaptive gene flow and in the evolution of reproductive isolation (Noor et al. 2001; Kirkpatrick and Barton 2006). Polygenic adaptation from standing genetic variation, the scenario characteristic of rapid environmental change and colonization events, represents precisely the ecological context in which indirect selection should favor elevated recombination: one defined by many segregating variants, high potential for Hill-Robertson interference, and thus the greatest opportunity for recombination to increase the efficacy of selection. Colonization events frequently involve founder effects, however, and our results suggest that such population contractions would substantially attenuate the benefit of recombination by depleting standing genetic variation. Beyond the goal of understanding variation in natural recombination rates, this result also has implications for the utility of recombination-rate modifiers in artificial selection scenarios (e.g., breeding). Given that domestication and subsequent breeding often involve strong demographic bottlenecks and founder effects (Hyten et al. 2006; Gaut et al. 2018), our results suggest that modification of recombination rate may not always accelerate breeding outcomes under such conditions. We may thus need to temper expectations about the potential benefits of genetic modification of recombination rates in agricultural species (Wijnker and Jong 2008; Mieulet et al. 2018).

Several features of our simulation design also bound the generality of our conclusions. Our simulations modeled a single chromosome with a fixed recombination landscape and did not allow recombination rates to evolve, precluding assessment of whether populations experiencing strong versus weak interference would evolve different rates, a question requiring an explicit modifier approach. Additionally, our LD analyses capture only alleles segregating at each measurement point, underrepresenting large-effect variants that are likely to have already fixed or been purged, and may therefore underestimate the true extent of Hill-Robertson interference during adaptation. Moreover, all three effect-size distributions we examined gave rise to polygenic architectures. While we found that genetic architecture strongly influenced adaptation rate, with few large-effect alleles accelerating adaptation more than threefold, the benefit of recombination did not differ across the architectures we modeled. Because the three gamma distributions shared a common mean effect size and differed only in their shape, even the most heavy-tailed distribution recruited hundreds of contributing loci, and the degree of realized polygenicity may not have varied enough to reveal an interaction with recombination rate. Architectures giving rise to truly oligogenic or monogenic adaptation might yield a different result. Empirical tests asking whether lineages with higher recombination rates show faster adaptation to novel environments would be a powerful complement to this simulation work, and the growing availability of high-quality recombination maps makes such comparisons increasingly feasible.

Understanding the evolutionary consequences of recombination rate variation is difficult in practice, because recombination rate covaries with features of genome organization and evolutionary history such that its independent effects are hard to isolate empirically. Forward-time simulations offer a complementary approach, allowing recombination’s effects to be examined in isolation across a controlled range of conditions. Our simulations, grounded in the biology of a well-characterized natural system and spanning a realistic range of genetic architectures and demographic scenarios, speak directly to recombination’s role during polygenic adaptation from standing genetic variation, the mode of adaptive evolution most relevant to rapid environmental change. The answer, at least within the parameter space we explored, is that more recombination consistently accelerates adaptation, and that this benefit operates primarily through the relief of Hill-Robertson interference. How much recombination accelerates adaptation, however, depends strongly on the demographic context. The populations most likely to benefit are those large enough for selection to act efficiently on the variation that recombination acts upon.

## Data availability

All code is available at https://github.com/samuk-lab/recomb-slim and all data is available at https://doi.org/10.6084/m9.figshare.33977680.

## Acknowledgements

This work was supported by the National Science Foundation Postdoctoral Research Fellowship in Biology awarded to M.A. (Award No. 2410227) and National Institutes of Health Grant No.

5R35GM154837-03 awarded to K.S.

## Author Contributions

M.A., E.E.A., and K.S. designed the study. M.A. designed and conducted the simulations and data analysis, and drafted the manuscript with critical input from all authors.

## References

Altenberg L, Feldman MW. 1987. Selection, generalized transmission and the evolution of modifier genes. I. The reduction principle. Genetics 117:559–572.

Barrett RDH, Schluter D. 2008. Adaptation from standing genetic variation. Trends Ecol Evol 23:38–44.

Barton NH. 1995. A general model for the evolution of recombination. Genet. Res. 65:123–144.

Barton NH, Charlesworth B. 1998. Why sex and recombination? Science 281:1986–1990.

Bates D, Mächler M, Bolker B, Walker S. 2015. Fitting linear mixed-effects models using lme4. J. Stat. Softw. 67.

Bell MA, Foster SA. 1994. The evolutionary biology of the threespine stickleback. Oxford, UK: Oxford University Press

Charlesworth B. 2009. Effective population size and patterns of molecular evolution and variation. Nat Rev Genet 10:195–205.

Cresko WA, McGuigan KL, Phillips PC, Postlethwait JH. 2007. Studies of threespine stickleback developmental evolution: progress and promise. Genetica 129:105–126.

Drury AL, Gout J-F, Dapper AL. 2023. Modeling recombination rate as a quantitative trait reveals new insight into selection in humans. Genome Biol. Evol. 15:evad132.

Feldman MW, Otto SP, Christiansen FB. 1996. Population genetic perspectives on the evolution of recombination. Annu. Rev. Genet. 30:261–295.

Felsenstein J. 1974. The evolutionary advantage of recombination. Genetics 78:737–756.

Fox J, Weisberg S, Price B. 2001. car: Companion to applied regression. CRAN: Contrib. Packag.

Gaut BS, Seymour DK, Liu Q, Zhou Y. 2018. Demography and its effects on genomic variation in crop domestication. Nat. Plants 4:512–520.

Haller BC, Ralph PL, Messer PW. 2025. SLiM 5: Eco-evolutionary simulations across multiple chromosomes and full genomes. Mol. Biol. Evol. 43:msaf313.

Hassold T, Hunt P. 2001. To err (meiotically) is human: the genesis of human aneuploidy. Nat. Rev. Genet. 2:280–291.

Hayward LK, Sella G. 2022. Polygenic adaptation after a sudden change in environment. eLife 11:e66697.

Hill WG, Robertson A. 1966. The effect of linkage on limits to artificial selection. Genet. Res. 8:269–294.

Höllinger I, Pennings PS, Hermisson J. 2019. Polygenic adaptation: From sweeps to subtle frequency shifts. PLoS Genet. 15:e1008035.

Hunt G, Bell MA, Travis MP. 2008. Evolution toward a new adaptive optimum: Phenotypic evolution in a fossil stickleback lineage. Evolution 62:700–710.

Hyten DL, Song Q, Zhu Y, Choi I-Y, Nelson RL, Costa JM, Specht JE, Shoemaker RC, Cregan PB. 2006. Impacts of genetic bottlenecks on soybean genome diversity. Proc. Natl. Acad. Sci. 103:16666–16671.

Jain K, Stephan W. 2017. Modes of rapid polygenic adaptation. Mol Biol Evol 34:3169–3175.

Jones GH, Franklin FCH. 2006. Meiotic crossing-over: Obligation and interference. Cell 126:246–248.

Kassambara A. 2019. rstatix: Pipe-friendly framework for basic statistical tests. CRAN: Contrib. Packag.

Kingman GAR, Vyas DN, Jones FC, Brady SD, Chen HI, Reid K, Milhaven M, Bertino TS, Aguirre WE, Heins DC, et al. 2021. Predicting future from past: The genomic basis of recurrent and rapid stickleback evolution. Sci. Adv. 7:eabg5285.

Kirkpatrick M, Barton N. 2006. Chromosome inversions, local adaptation and speciation. Genetics 173:419–434.

Kuznetsova A, Brockhoff PB, Christensen RHB. 2017. lmerTest package: Tests in linear mixed effects models. J. Stat. Softw. 82.

Lenth RV, Piaskowski J. 2017. emmeans: Estimated marginal means, aka least-squares means. CRAN: Contrib. Packag.

Lin M, Chakraborty S, Amorim CEG, Nigenda-Morales SF, Beichman AC, Nuñez-Valencia PG, Mah J, Robinson JA, Kyriazis CC, Huber CD, et al. 2025. The distribution of fitness effects varies phylogenetically across animals. bioRxiv:2025.05.13.653358.

Mieulet D, Aubert G, Bres C, Klein A, Droc G, Vieille E, Rond-Coissieux C, Sanchez M, Dalmais M, Mauxion J-P, et al. 2018. Unleashing meiotic crossovers in crops. Nat. Plants 4:1010–1016.

Montgomery EA, Huang SM, Langley CH, Judd BH. 1991. Chromosome rearrangement by ectopic recombination in Drosophila melanogaster: genome structure and evolution. Genetics 129:1085–1098.

Muller HJ. 1964. The relation of recombination to mutational advance. Mutat Res Fundam Mol Mech Mutagen 1:2–9.

Nickel J, Foote AD. 2026. Hitchhiking of deleterious mutations within chromosomal inversions. Trends Ecol Evol.

Noor MAF, Grams KL, Bertucci LA, Reiland J. 2001. Chromosomal inversions and the reproductive isolation of species. Proc National Acad Sci 98:12084–12088.

Orr HA, Betancourt AJ. 2001. Haldane’s sieve and adaptation from the standing genetic variation. Genetics 157:875–884.

Otto SP, Barton NH. 1997. The evolution of recombination: removing the limits to natural selection. Genetics 147:879–906.

Otto SP, Barton NH. 2001. Selection for recombination in small populations. Evolution 55:1921–1931.

Otto SP, Lenormand T. 2002. Resolving the paradox of sex and recombination. Nat Rev Genet 3:252–261.

Reid K, Bell MA, Veeramah KR. 2021. Threespine stickleback: A model system for evolutionary genomics. Annu. Rev. Genom. Hum. Genet. 22:1–27.

Ripley B, Venables B. 2009. MASS: Support functions and datasets for venables and Ripley’s MASS. CRAN: Contrib. Packag.

Ritz KR, Noor MAF, Singh ND. 2017. Variation in recombination rate: adaptive or not? Trends Genet 33:364–374.

Rockman MV. 2012. The QTN program and the alleles that matter for evolution: all that’s gold does not glitter. Evolution 66:1–17.

Roesti M, Gilbert KJ, Samuk K. 2022. Chromosomal inversions can limit adaptation to new environments. Mol. Ecol. 31:4435–4439.

Roesti M, Moser D, Berner D. 2013. Recombination in the threespine stickleback genome—patterns and consequences. Mol. Ecol. 22:3014–3027.

Samuk K, Owens GL, Delmore KE, Miller SE, Rennison DJ, Schluter D. 2017. Gene flow and selection interact to promote adaptive divergence in regions of low recombination. Mol Ecol 26:4378–4390.

Scholz F, Zhu A. 2025. kSamples: K-sample rank tests and their combinations. CRAN: Contrib. Packag. [Internet]. Available from: https://CRAN.R-project.org/package=kSamples

Shanfelter AF, Archambeault SL, White MA. 2019. Divergent fine-scale recombination landscapes between a freshwater and marine population of threespine stickleback fish. Genome Biol. Evol. 11:1573–1585.

Stapley J, Feulner PGD, Johnston SE, Santure AW, Smadja CM. 2017. Variation in recombination frequency and distribution across eukaryotes: patterns and processes. Philosophical Transactions Royal Soc B Biological Sci 372:20160455.

Team RC. 2024. R: A language and environment for statistical computing. Available from: https://www.R-project.org/

Thompson JN. 1997. Evaluating the dynamics of coevolution among geographically structured populations. Ecology 78:1619–1623.

Tigano A, Friesen VL. 2016. Genomics of local adaptation with gene flow. Mol Ecol 25:2144–2164.

Wang H, Zhang C, Reid K, Merilä J. 2026. Recombination rate and efficiency of linked selection in small and large stickleback populations. bioRxiv:2026.03.18.712813.

Wickham H, Averick M, Bryan J, Chang W, McGowan L, François R, Grolemund G, Hayes A, Henry L, Hester J, et al. 2019. Welcome to the Tidyverse. J. Open Source Softw. 4:1686.

Wijnker E, Jong H de. 2008. Managing meiotic recombination in plant breeding. Trends Plant Sci. 13:640–646.

Wilson BA, Petrov DA, Messer PW. 2014. Soft selective sweeps in complex demographic scenarios. Genetics 198:669–684.

Yeaman S, Whitlock MC. 2011. The genetic architecture of adaptation under migration-selection balance. Evolution 65:1897–1911.

Zhang C, Reid K, Schierup MH, Wang H, Candolin U, Merilä J. 2025. Rate of de novo mutations in the three-spined stickleback. Heredity 134:387–395.

